# Microrobot and Cellbot Locomotion in Confined Hydrogels and Cardiac Tissue Surfaces

**DOI:** 10.64898/2026.10.19.760203

**Authors:** Fatma Ceren Kirmizitas, Sudipta Mallick, Subrahmanyam Cherukumilli, Zion D. Lee, Yuwen Zhao, Aditya Dutta, Yaling Liu, Sambeeta Das

**Affiliations:** Department of Mechanical Engineering University of Delaware, Newark, DE 19716 USA; Department of Animal and Food Sciences, University of Delaware, Newark, DE 19716 USA; Center for Organoid and Organ on Chip, West China Hospital, Sichuan University, Chengdu, Sichuan 610041 China; Center for High Altitude Medicine, West China Hospital, Sichuan University, Chengdu, Sichuan 610041 China; Lehigh University, Bethlehem, PA 18015 USA

**Keywords:** Microrobots, manipulation, cardiac tissue, hydrogels

## Abstract

Biohybrid cellbots and microrobots are a promising platform for targeted drug delivery, minimally invasive surgery, and tissue engineering. Here, we present a comparative analysis of the locomotion dynamics across synthetic substrates and ex vivo cardiac tissue to help bridge the gap presented by translating locomotion dynamics from idealized environments to biological interfaces. By subjecting the microrobots and cellbots to different substrates (specifically, cell culture dish, hydrogel, and ex vivo tissue) and applying rotating magnetic fields at varying frequencies, we characterized how surface boundary interactions and driving frequency dictate translation speed. We reveal that particles moving on tissue exhibit restricted movement based on trajectory and tissue fiber orientation. Finally, we discuss that while cellular integration introduces significant adhesive drag that reduces translation speeds, it simultaneously offers a unique clinical advantage. This study highlights the potential of balancing magnetic driving parameters against biological environments to optimize for various treatment timelines and efficiency across diverse biomedical applications.

## I. Introduction

Microrobots capable of controlled locomotion in complex fluidic and biological environments have emerged as a promising platform for applications in targeted drug delivery, minimally invasive surgery, and tissue engineering. Microrobots are particularly applicable in narrow, tortuous, and mechanically heterogeneous biological spaces where conventional treatments are less effective [1]. The design and applications of microscale robots are constrained by the physics of motion at the microscale. Since inertia is negligible and the Reynolds number is low, motion at the microscale is governed by Stokes flow. Purcell’s scallop theorem outlines that non-reciprocal strokes are needed to generate net translation [2], [3]. In this regime, locomotion requires broken time-reversal symmetry, whether through helical propulsion, peristaltic shape change, surface rolling, or other nonreciprocal strategies that exploit geometry, compliance, or boundary interactions. Thus, microrobot propulsion is never purely a property of the actuator; it is always a property of the actuator coupled to its environment.

Recently, the hydrodynamic interactions of microrobots and their surrounding surfaces have gained interest as they are an important factor. Physiologically relevant vessel-like structures [4], in vitro cell dense environments [5], and ex vivo bovine oviduct tissue [6] have been studied to demonstrate the various microrobot movements. These studies are extremely valuable for understanding the realistic conditions and nature of targeted biological environments that present unique challenges. Biohybrid microrobots extend this logic by embedding living cells as actuators, sensing units, or both [7]. Their appeal is that biological components bring intrinsic motility, adaptive response, and biochemical functionality that are difficult to reproduce synthetically. Previously, our group has presented the use of cellbots to demonstrate autonomous control and cellular applications [8], [9]. Cardiomyocyte-driven constructs, in particular, have been explored for their ability to generate rhythmic forces and respond dynamically to biochemical cues, enabling functionalities that are difficult to replicate with purely synthetic systems [10], [11], [12]. Cells have also been combined with artificial carriers to achieve multimodal control and cargo delivery in vivo [13]. Despite substantial progress in both synthetic microrobotics and biohybrid actuation, most studies have focused on locomotion within either engineered or biological environments in isolation. In contrast, many real-world biomedical applications require navigation across heterogeneous interfaces, such as transitions between synthetic scaffolds (e.g., hydrogels) and living tissues. Hydrogels are widely used as model extracellular matrices due to their tunable stiffness, porosity, and biochemical composition. However, living tissues such as cardiac muscle have fundamentally different conditions, such as dynamic mechanical activity, heterogeneous surface topography, and active force generation, all of which can significantly alter locomotion mechanisms [14]. Here, we study the locomotion and interaction dynamics of synthetic microrobots and biohybrid cellbots in two representative environments: confined hydrogel channels and ex vivo cardiac tissue surfaces. By directly comparing control strategies and substrate interactions across these platforms, this work serves as a proof-of-concept framework for understanding cross-interface locomotion and provides design guidelines for next-generation microrobotic systems (Fig. 1). In subsequent studies, we plan to study the governing physical and biological mechanisms underlying mobility in heterogeneous media using this system.

**Figure 1.**
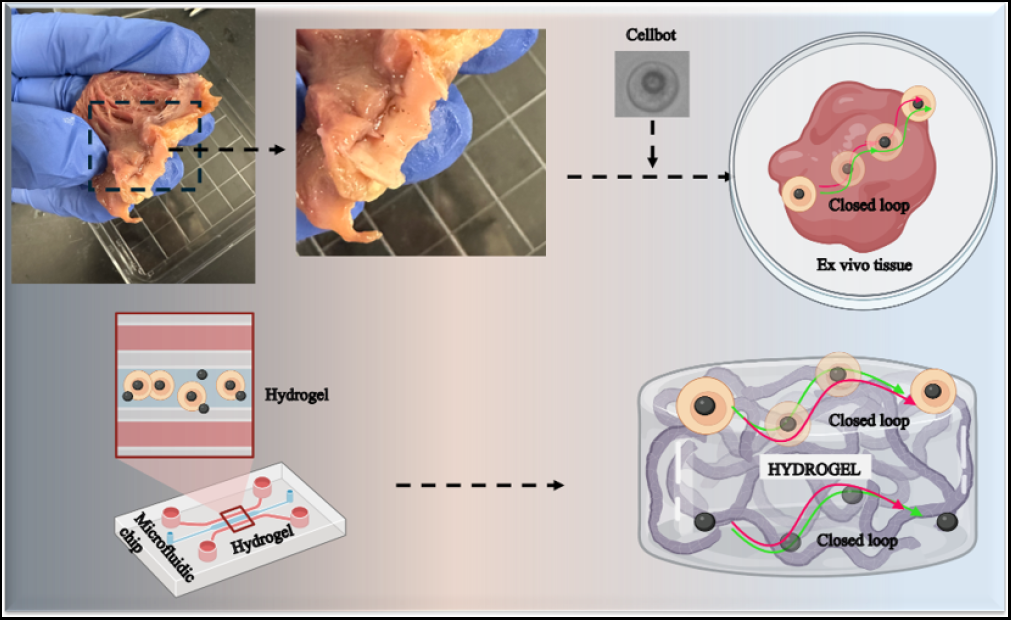
Schematic of the experimental setup. Cellbots and microrobots are manipulated in hydrogel channels and on cardiac tissue surface with external magnetic system.

## II. Material and Methods

### A. Fabrication of microrobots and operation

The microrobots were fabricated using commercially available 20 *µ*m diameter silica spheres with amine functionality (Nanocs, Cat.No.Si20u-AM-1) and 4.8 *µ*m (CliniSciences, Cat.No.SSD50003) diameter silica particles. The silica spheres were pipetted onto a glass substrate to form a uniform layer. Subsequently, the spheres are coated with 100 *µ*m of nickel (Ni) layer using the physical vapor deposition technique to impart magnetic properties. The deposition was performed perpendicular to the substrate, resulting in a hemispherical coating on the microrobots. The magnetic moment of the half-coated microrobots tends to orient normally to the surface.

### B. Experimental setup

A schematic representation of the experimental design is shown in Fig. 2. The experimental setup consists of a 3D Helmholtz coil system comprised of 6 pairs of electromagnetic coils to generate a uniform magnetic field of 5mT at the center. The coil system is mounted on an Axiovert 200 inverted microscope. The electromagnetic coil system is powered by a DC power supply. An arduino based controller system consisted of H-bridges and PWM Drivers to regulate the input current load and direction to the coils, is implemented for computational abilities. A custom Python code is used to modulate the current sent to each coil for precise control.

**Figure 2.**
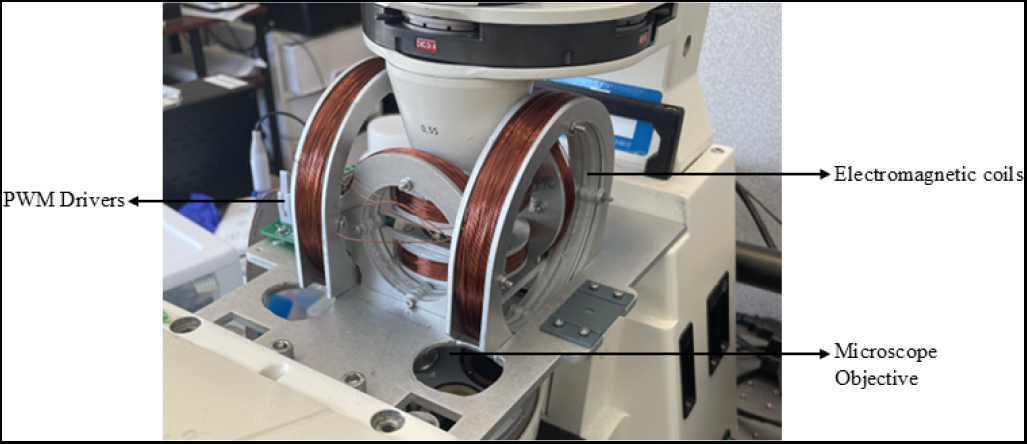
Experimental Setup showing the electromagnetic coil system.

### C. Cell culture and fabrication of cellbots

Chinese Hamster Ovary (CHO) cells were used to demonstrate cellular mobility. CHO cells were kindly provided by our collaborator Dr. Ron Weiss (Massachusetts Institute of Technology, Massachusetts, USA). All cell culture components were purchased from GIBCO, USA. CHO cells were cultured in Dulbecco’s Modified Essential Medium F-12 (DMEM/F-12) media supplemented with 10% Fetal Bovine Serum (FBS) and 1% penicillin-streptomycin in a standard cell culture incubator with 5% *CO*_2_ at 37 °C. All experiments were performed after the third passage, and before each experiment, cells were washed with Dulbecco’s phosphate buffer saline (DPBS) and detached with TrypLE Express from the cell culture dish. Our group has previously used CHO cells to fabricate the cellbots for various cellular applications. It has been shown that the microrobot internalization does have a minimal effect on cell viability and proliferation, as well as the external magnetic control system on CHO cells [9]. Briefly, to fabricate cellbots, CHO cells were seeded in a 35 *mm* cell culture dish at a density of 1*×*10^5^ cells*·*mL^*−*1^. To seed the cells on a dish, trypan blue staining was performed. 10 *µ*l cell suspension was mixed with 0.4% trypan blue in a 1:1 (v/v) ratio, and cells were counted on a cell counter (Nexcelom Cellometer Vision Trio Cell Profiler, USA). 4.8 *µ*m Nickel-coated microrobots were scratched off the glass slide, sonicated, and collected by using a handheld magnet to obtain a well-dispersed microrobot solution. Cells were incubated under the same standard cell culture conditions for 24 hours, and the next day, cells were detached and washed to obtain cellbots. Cellbot preparation steps and microscope image are shown in Fig. 3

**Figure 3.**
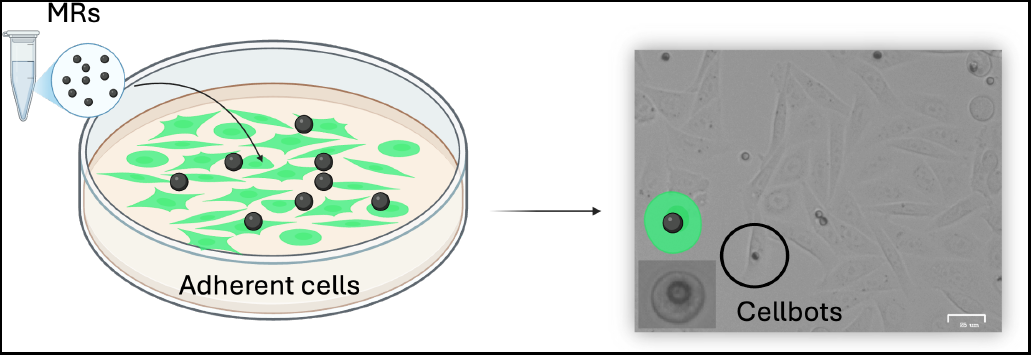
Cellbot preparation and optical microscope image of cellbots. CHO cells were incubated with 4.8 *µ*m size microrobots overnight to obtain cellbots.

### D. Magnetic control

The motion of the microrobots is primarily due to the magnetic torque generated by the rotating magnetic fields. The equation of motion is given by

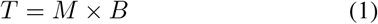

where M is the Magnetic moment, and B is the Magnetic field. The translation motion of the microrobot in x and y directions on a 2D plane could be achieved by applying rotating magnetic fields either in the xz or yz planes, respectively. The speed of the microrobots is linearly dependent on the magnetic frequency applied. The motion is also affected by the various forces acting between the microrobot and substrate surface, such as Van Der Waals and hydrodynamic forces. The rotating magnetic fields were applied according to the following equations,

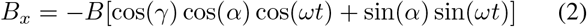

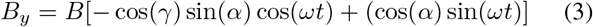

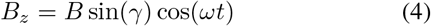

Where *γ* is the azimuthal angle from the z axis, *α* is the polar angle from the x axis, B is the magnetic field magnitude, and *ω* is the frequency that controls the speed of the rolling microrobot. An azimuthal angle of 90^*°*^ was used in all of the experiments presented here.

### E. Ex vivo tissue preparation

All procedures involving animals, including acquisition, housing, care, and euthanasia, were conducted in accordance with protocols approved by the University of Delaware Animal Care and Use Committee (IACUC). Animals were fed once daily and had unrestricted access to water. From a flock of 150 two-year-old specific pathogen-free (SPF) White Leghorn laying hens obtained from Charles River Laboratories (Norwich, CT, USA), two individuals were randomly chosen and euthanized via cervical dislocation. After confirming death, the hens were dissected, and their hearts were perfused with 1X phosphate-buffered saline (PBS) to remove residual blood. The hearts were then excised and transported in sterile specimen containers (McKesson, TX, USA) containing 1X PBS for subsequent tissue processing. Cardiac tissue was sectioned into flat 20 *µ*m slices using a cryostat (Avantik QS12, Pine Brook, NJ) and mounted onto SuperFrost Plus slides (Fisher Scientific, USA). The prepared tissue sections were stored at -20 °C until further use.

### F. Microfluidic hydrogel channel preparation

To test the navigation efficacy of the microrobots and cellbots, we used a microfluidic hydrogel device. The microfluidic channel was designed in AutoCAD and fabricated using standard soft lithography with SU-8 2150 (Kayaku Advanced Materials) to achieve a chamber height of 800 µm, as mentioned previously [15], [16], [17]. The polydimethylsiloxane (PDMS, SYLGARD 184, Dow Corning Corporation) was used to mold through the SU8 mold and was solidified at 50 °C overnight, followed by plasma bonding to the 1-mm-thick glass slide (Amscope). All the loading ports were punched with 1.5 mm and 3 mm biopsy punches (World Precision Instruments), respectively. To generate the hydrogel channel, fibrinogen and bovine thrombin (Sigma) were mixed and injected into the chamber at final concentrations of 10 mg/mL fibrinogen and 5 U/mL thrombin, forming a fibrin gel. To create a hollow hydrogel channel, the PDMS rod was molded using a 27-gauge needle and then extruded after the fibrin gel had crosslinked. The dimensions of the channel is 210 *µ*m in diameter.

## III. Results and Discussion

### A. Motion on ex vivo cardiac tissue

Initial experiments are performed to study the feasibility of the microrobots’ motion on the ex vivo cardiac tissue surface. To study the microrobot locomotion on a biological surface, a cardiac tissue slice attached to a glass slide was used as the substrate. Firstly, 20 *µ*m microrobots were added onto the substrate using DI water. Once the rotating magnetic fields are applied, the microrobots can roll along the x and y directions depending on the direction of the applied field using a wireless controller as in Fig. 4. It can be seen that the microrobot was able to move in all four directions. The magnitude of the magnetic field can be controlled by the amount of force the joystick is pushed. The speed of the microrobots is dependent on the frequency applied. Besides, the motion of the microrobots on the surface is affected by the friction and the hydrodynamic drag forces acting between the microrobot and the substrate surface.

**Figure 4.**
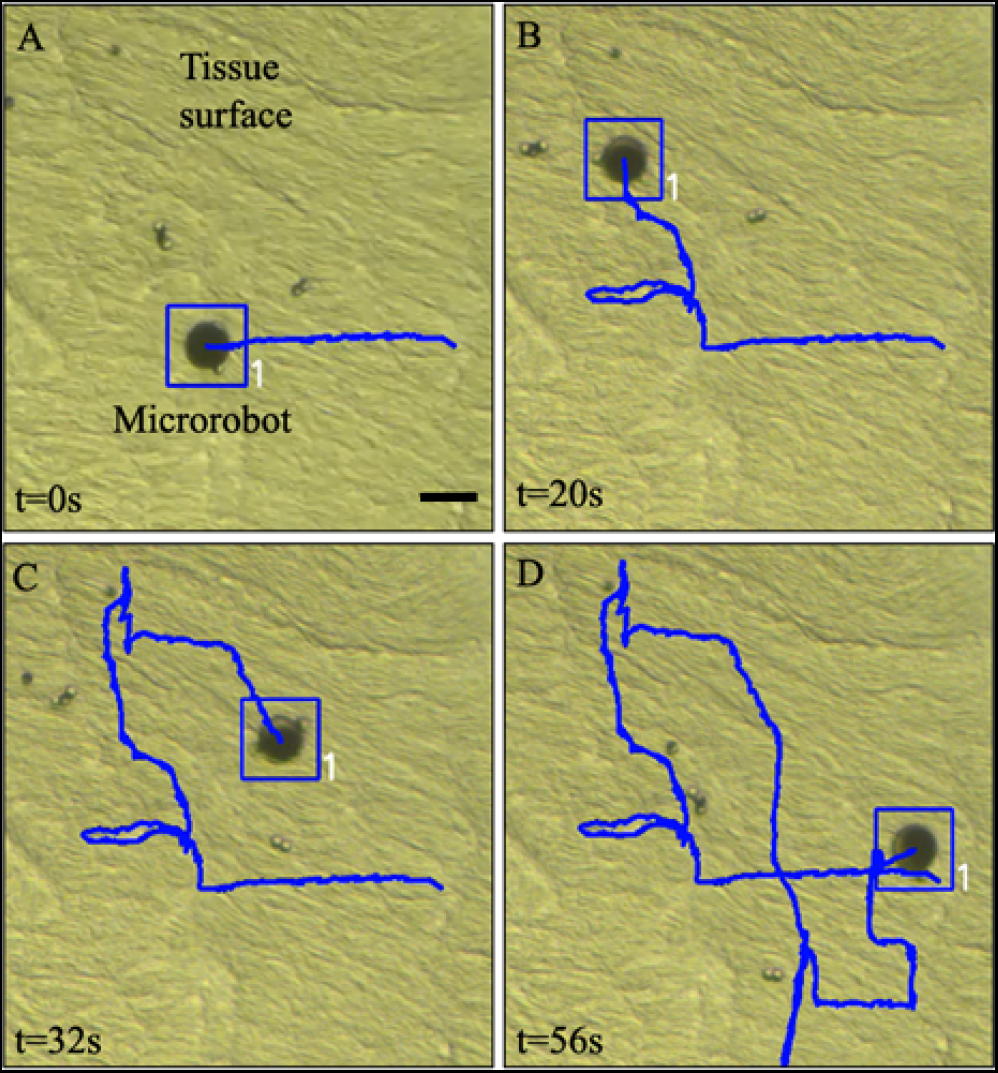
Open loop control of microrobot onto the surface of cardiac tissue over time (A)-(D). Scale bar is 20 *µ*m.

In addition, the navigation of 4.8 *µ*m Nickel-coated microrobots was also determined on a cardiac tissue slice, as shown in Fig. 5 and actuated using an applied magnetic field. The ex vivo tissue slide was approximately 20 *µ*m in thickness. The microrobot could move onto the tissue surface in 17 seconds in total. In addition, fluid-induced drag also contributed to the resistance of the microrobot’s locomotion. The cellular architecture and alignment are critical for tissue organizations, especially for cardiac tissue metabolism and function [18].

**Figure 5.**
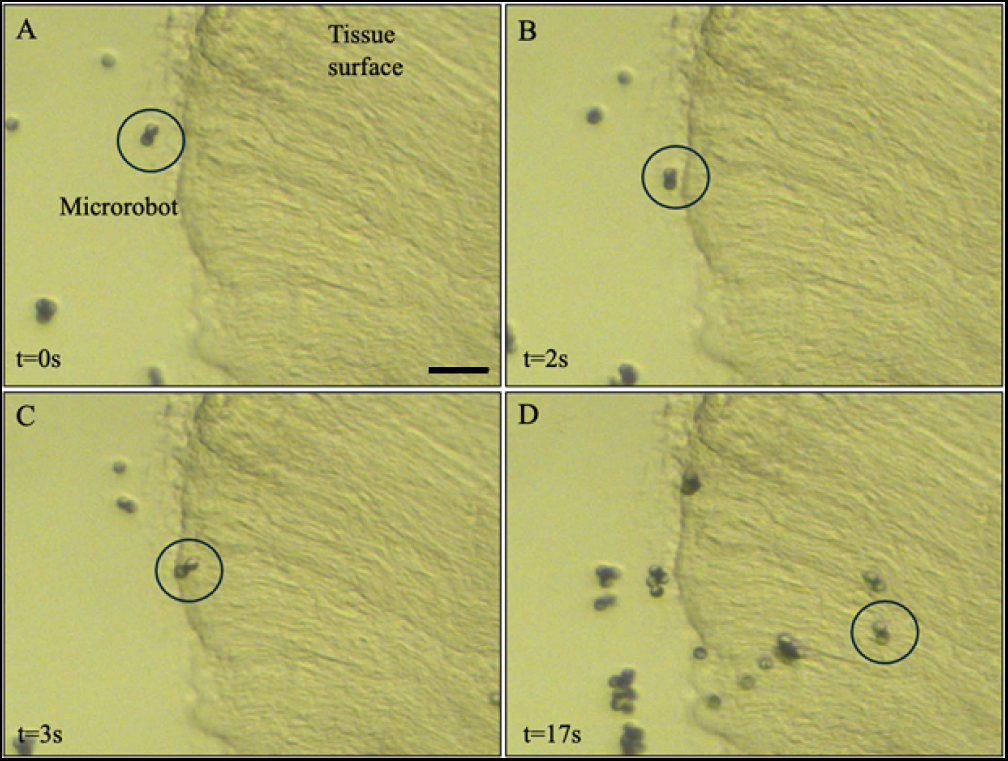
4.8*µ*m microrobot locomotion on the ex vivo cardiac tissue surface (A)-(D). Scale bar is 20 um.

A successful in vitro and ex vivo model should be able to demonstrate the cell behavior in a spatiotemporal manner. To achieve this, we evaluated the controlled navigation of cellbots on the ex vivo cardiac tissue since the tissue slide preserves the grooves and uneven surface topography of the heart tissue. Subsequently, we used the biohybrid robots to determine the efficacy and potentials onto the tissue. Cellbots were prepared as previously described and were placed on a cardiac tissue slice, as shown in Fig. 6. A single cellbot was placed on the petri dish and moved towards the cardiac tissue with the external magnetic field. The cells are now responsive to the magnetic field following the uptake of the 4.8 *µ*m microrobots and can be moved on the tissue surface in approximately 10 seconds. The cellbot locomotion on the tissue was slower than that of the microrobots. This additional speed reduction was attributed to adhesion between the cellbots and the tissue surface.

**Figure 6.**
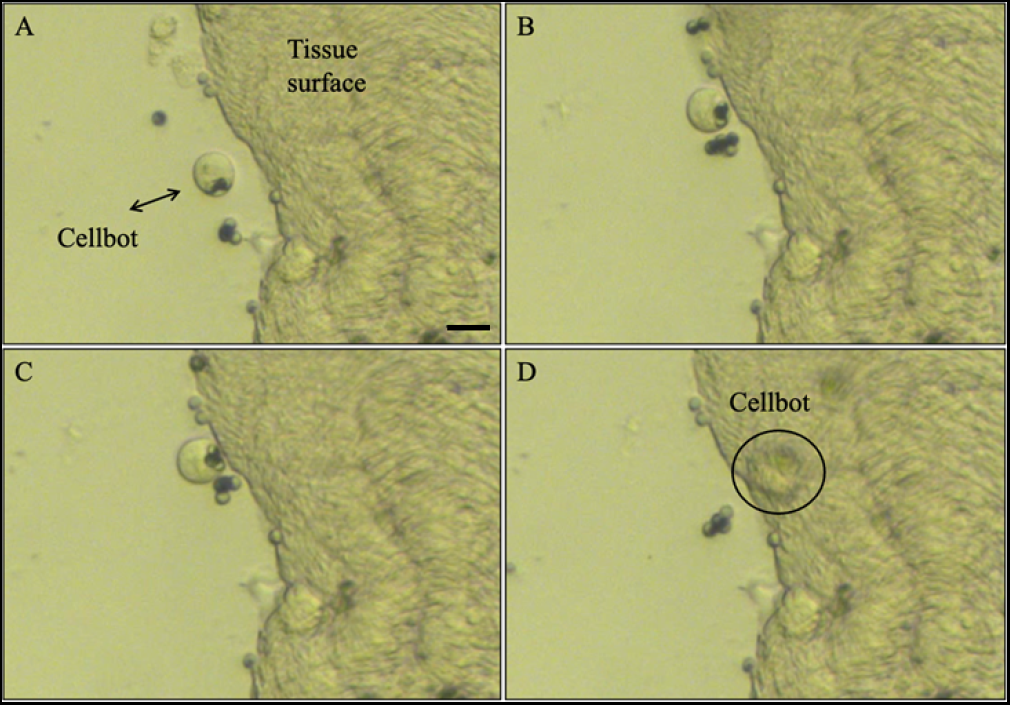
CHO cellbot movement onto the surface of cardiac tissue (A)-(D). Scale bar is 10 *µ*m.

The targeted delivery of cells is a critical effect of the cell delivery in a specific alignment. Previously, several groups have shown the cardiomyocyte cultures on 2D environments without targeted alignment or delivery strategies; yet, the possibility of cells growing in unexpected directions can lead to cellular contraction and mechanical stress [19], [20]. To overcome the unexpected deliveries and subsequent challenges, a custom-made closed-loop control system is implemented to demonstrate the feasibility of the microrobot’s and cellbot’s motion by following a predefined trajectory from a starting point to the ending point. A complex trajectory of “SMT” is defined as shown in Fig. 7C, on a cell-dense hydrogel surface by the user. The algorithm iteratively computes the difference between the current node’s position and the desired position, then moves on to the next node if the error is less than a predefined threshold. When the inaccuracy exceeds the threshold, the algorithm calculates the magnetic field vector required to direct the microrobot to the intended position, taking into account the characteristics of the equations mentioned in 2, 3, 4 [21]. The microrobots followed the predefined trajectory on the hydrogel surface with a minimal overshoot and completed the path in 15 seconds with an average speed of 30 um/s. Previously, hydrogel was used to mimic the natural structure of the tissue, while cells were aligned inside in the presence of the magnetic field via the magnetic nanoparticle internalization [22]. Cells were responsive to the magnetic field, and could be aligned inside the gel within 30 minutes via permanent magnets; however, this alignment is missing the autonomous fashion. In this work, we use a sophisticated external magnetic control system to control cellbots and microrobots to successfully test the efficacy of delivering methods on the ex vivo surfaces. It should be noted that the microrobot locomotion on the tissue surface was slower than on the hydrogel surface because the tissue’s uneven topography increased stiction and friction. In our previous work, we have already shown the cellular delivery and patterning via both open-loop and closedloop control, and we plan to fabricate novel cardiomyocyte-based cellbots to evaluate their potential in the cardiovascular tissue engineering applications [23], [24].

**Figure 7.**
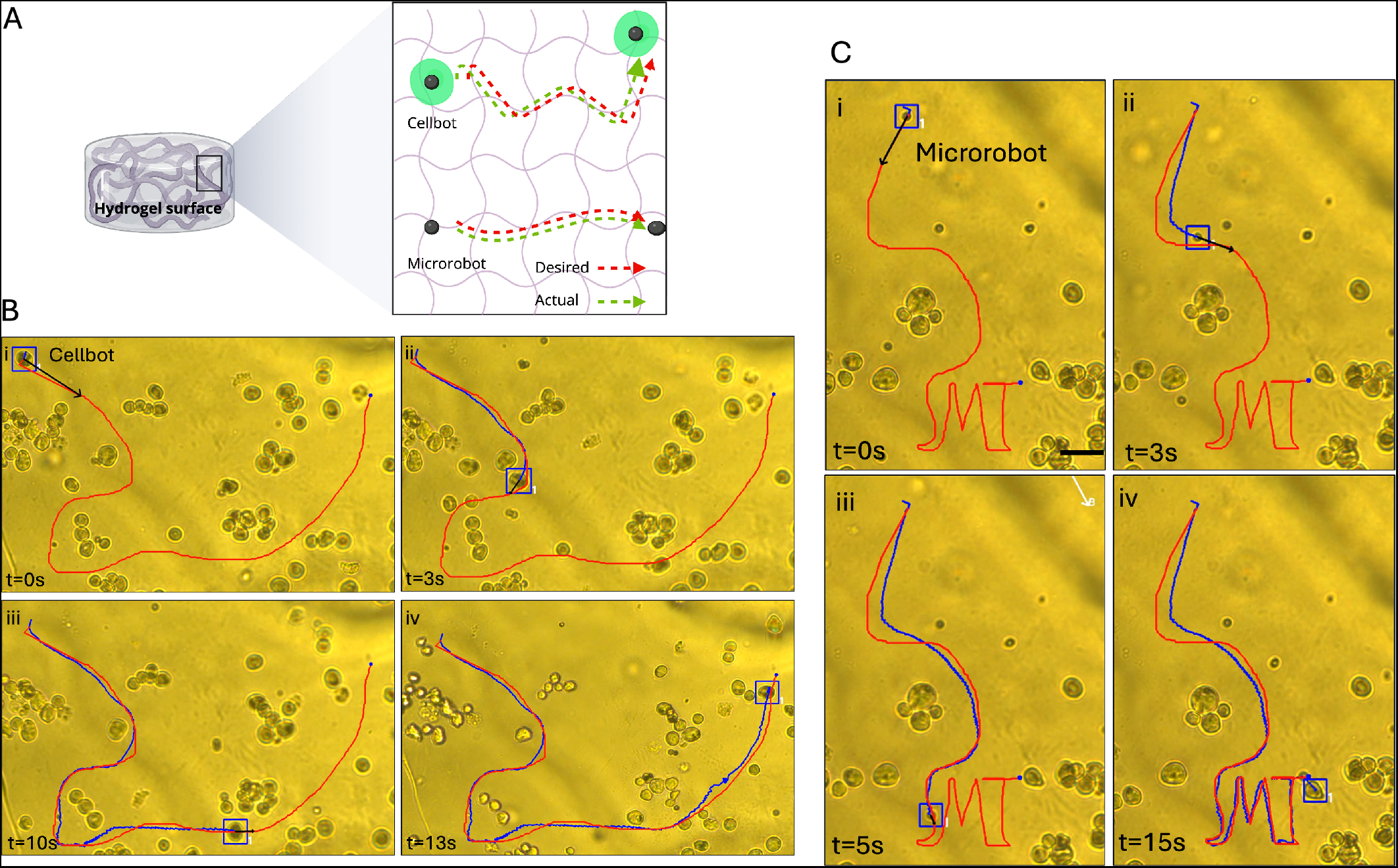
Microrobot and cellbot locomotion via closed-loop control system on the hydrogel surface over time. A) Schematic of the navigation of the cellbot and microrobots. Series of images of closed-loop control of cellbots (Bi-iv) and microrobots (Ci-iv) on a hydrogel surface from the starting point to the end point over 13 and 15 seconds, respectively. Scale bar is 20 *µ*m.

### B. Comparative analysis

The speed of the microrobots and cellbots under variable magnetic frequency has been analyzed in this section. A microscope glass slide served as a control to establish baseline speed profiles for the particles. Ni-coated microrobots and cellbots (measuring approximately 20 *µ*m in diameter) were tested on both the glass substrate and on the ex vivo cardiac tissue, with results shown in Fig. 8. Locomotion of the microrobots showed significantly higher locomotion speeds to that of cellbots over the range of tested frequencies.

**Figure 8.**
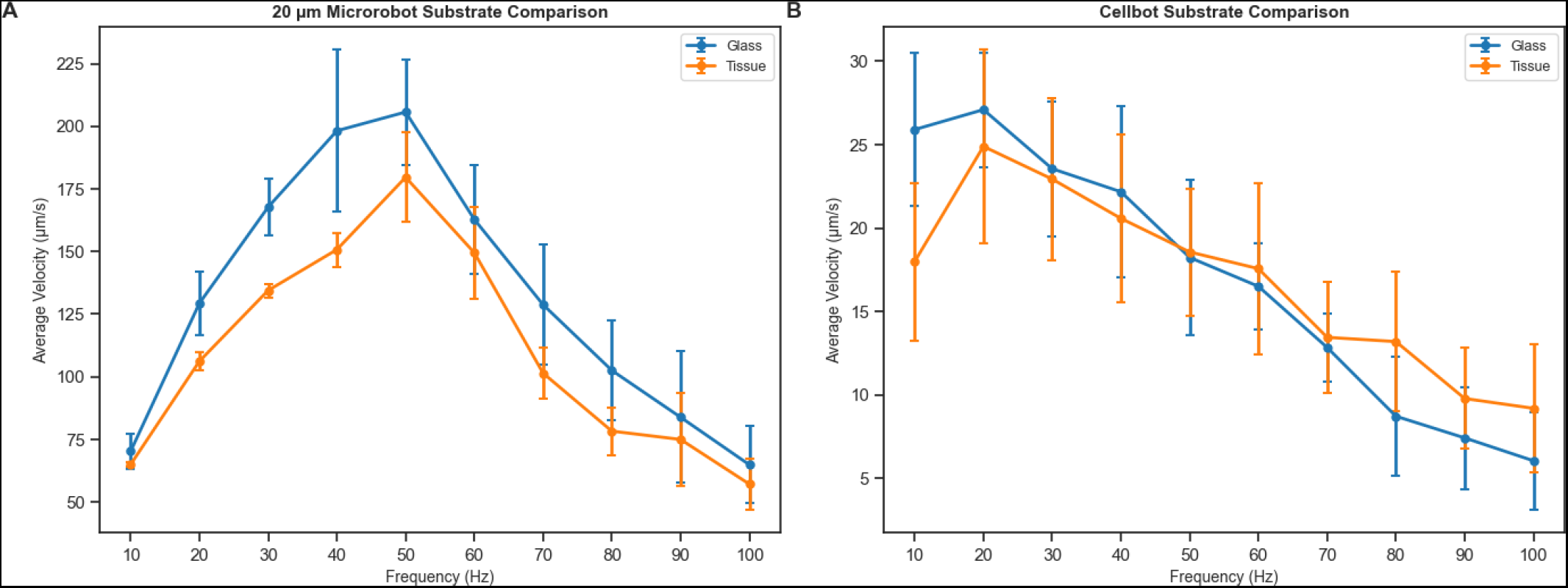
Comparison between the speed [*µ*m/s] vs Frequency [Hz] of 20 *µ*m microrobots and cellbots on glass and tissue substrates.

The speed of the Ni-coated 20 *µ*m microrobots, across both substrates reach a maximum average speed of 179.6 and 205.7 *µ*m/s at approximately 50 Hz, across tissue and glass, respectively. Locomotion on cardiac tissue yielded slightly lower speeds than on the dish, however, this marginal reduction in translation speed is accompanied by lower variance in speed.

On the other hand, cellbots exhibited an overall negative speed-versus-frequency trend and achieved a maximum average translation speed of 24.9 and 27.1 *µ*m/s at approximately 20 Hz across tissue and glass, respectively. Interestingly, at higher frequencies (50 - 100 Hz), cellbots traversing cardiac tissue maintained a higher speed than on glass. Although this variance in translation speed across the tested range was minimal, the cellbots stopped showing effective rolling behavior above 40 Hz, failing to traverse at least one body length per second.

## IV. Conclusion

In this paper, we systematically characterized and compared the locomotion dynamics and surface interactions of synthetic nickel-coated microrobots and biohybrid cellbots across synthetic and biological substrates. Our experimental results revealed several critical insights that can help bridge the gap between idealized laboratory environments and clinical applications.

The use of CHO cells to form cellbots fundamentally alters the particle dynamics under rotating magnetic fields. Cellbots exhibit a stabilized speed profile across substrates due to the host cell. This inherent stability of cellbots comes at the cost of translation speed, which is drastically lower than a microrobot due to decreased magnetic torque, increased viscous resistance, and active biological adhesion. Lastly, our evaluation of the 20 *µ*m microrobots has shown that microtopography acts as a physical barrier where grooves and tissue fibers impose resistance to trajectory-dependent translation. Future potential applications could be more focused on cardiac cell-focused biohybrid microrobots to be used in the cardiovascular tissue engineering field. The controlled navigation in confined and complex environments could be a great advantage in cell patterning, alignment, and biological substrate deliveries to the target site.

## V. Acknowledgments and Funding

Figures were created with BioRender.com using an institutional license sponsored by the University of Delaware Office for Research. This research was funded by University of Delaware GUR award.

## VI. Data Availability

Data are available upon reasonable request.

## VII. Authors Contribution

F.C.K.: Conceptualization, Investigation, Visualization, Writing – original draft, Writing - review & editing. S.M.: Conceptualization, Investigation. S.C.: Investigation, Methodology, Writing – original draft, Writing - review & editing. Z.D.L.: Investigation, Visualization, Writing – original draft, Writing - review & editing. Y.Z.: Investigation, Writing - original draft. A.D.: Investigation. Y.L.: Investigation. S.D.: Funding acquisition, Supervision, Writing-review and editing, Resources.

